# A naturally-occurring dominant-negative competitor of Keap1 against its inhibition of Nrf2

**DOI:** 10.1101/286997

**Authors:** Lu Qiu, Meng Wang, Yuping Zhu, Yuancai Xiang, Yiguo Zhang

## Abstract

Transcription factor Nrf2 is a master regulator of antioxidant and/or electrophile response elements (AREs/EpREs)driven genes involved in homeostasis, detoxification and adaptation to various stresses. The cytoprotective activity of Nrf2, though being oppositely involved in both cancer prevention and progression, is critically controlled by Keap1 (Kelch-like ECH-associated protein 1) as an adaptor subunit of Cullin 3-based E3 ubiquitin ligase, that is a key sensor for oxidative and electrophilic stresses. Now, we first report a novel naturally-occurring mutant of Keap1, designated Keap1^ΔC^, which lacks most of its C-terminal Nrf2-interacting domain essential for inhibition of the CNC-bZIP factor. This mutant Keap1^ΔC^ is yielded by translation from an alternatively mRNA-spliced variant lacking the fourth and fifth exons, but their coding sequences are retained in the wild-type *Keap1* locus (with no genomic deletions). Although this variant was found primarily in the human highly-metastatic hepatoma (MHCC97H) cells, it was widely expressed at very lower levels in all other cell lines examined. No matter whether Keap1^ΔC^ retains less or no ability to inhibit Nrf2, it functions as a dominant-negative competitor of Keap1 against its inhibition of Nrf2-target genes. This is due to its antagonist effect on Keap1-mediated turnover of Nrf2 protein.

## INTRODUCTION

The gene encoding Nrf2 (nuclear factor-erythroid 2-related factor 2) was first cloned in 1994 and later identified as one of the most important members of the cap’n’collar (CNC) basic-region leucine zipper (bZIP) family [1, 2]. Most of studies about the CNC-bZIP family have been focused disproportionately on Nrf2 [3], because it is generally accepted as a master regulator of antioxidant and/or electrophile response elements (AREs/EpREs)-driven genes, which are involved in important homeostasis, detoxification and adaptive cytoprotection against various stresses [4, 5]. The activity of Nrf2 is executed through its functional heterodimer with its partner small Maf or other bZIP proteins, insomuch as they enable for binding to downstream target genes. Such transcriptional activity of Nrf2 is also negatively regulated by Keap1 (Kelch-like ECH-associated protein 1), because their direct interaction enables this CNC-bZIP protein to be sequestered within the cytoplasmic subcellular compartments, where it is targeted for its ubiquitination by Keap1, acting as an adaptor subunit of Cullin 3-based E3 ubiquitin ligase, as a result leading to its protein turnover through ubiquitin-mediated proteasomal degradation [6, 7]. However, upon stimulation of Keap1 as a key sensor for oxidative and electrophilic stresses, Nrf2 is dissociated from Keap1-sequestered confinements to be translocated into the nucleus before transactivating AREs/EpREs-battery genes, such as those encoding heme oxygenase-1 (HO-1), glutamate-cysteine ligase modifier subunit (GCLM), and NAD(P)H:quinone dehydrogenase 1 (NQO1) [8–10].

In fact, wild-type Keap1 was first identified to act as an Nrf2-specific inhibitor until 1999 [11, 12], and shares highly evolutionary conservation with the *Drosophila* Kelch protein, which is essential for the formation of actin-rich intracellular bridges termed ring canals [13]. Further studies have demonstrated that the negative regulation of Nrf2 by Keap1 is exerted through direct interaction of its C-terminal six double glycine-repeat (DGR)-adjoining domain with the Neh2 region of this CNC-bZIP protein, controlling the protein turnover [14, 15]. Structural studies have also revealed that only a functional homodimer of Keap1 with each other’s BTB domains is necessary for direct binding to the ETGE and DLG motifs within the Neh2 domain of Nrf2[15], which acts as a degron targeting the CNC-bZIP protein to ubiquitin-mediated proteasomal degradation pathways.

Whilst global knockout of Nrf2 in the mouse does not alter normal physiological development and growth with no spontaneous pathological phenotypes, all *Nrf2*^-/-^ animals are more susceptible than wild-type mice to chemical cytotoxicity [16]. By contrast, global knockout of its inhibitor Keap1 causes a constructive enhancement in both protein stability and transactivation activity of Nrf2, but leading to death of all *Keap1*^*-/-*^-deficient mice within three weeks after birth [17, 18]. This lethality is principally attributed to malnutrition caused by severe hyperkeratosis within the upper digestive tract (i.e. esophagus and forestomach). Further analysis of *Keap1:Nrf2* double knockout mice has confirmed that the recognizable phenotype of *Keap1*^*-/-*^-deficient mice is attributable to the constitutive activation of Nrf2. To avoid such death of murine pups before weaning, conditional hepatocyte-specific *Keap1*^*-/-*^ mice were generated, as an expected result of viable animals with no apparent abnormalities, but a nuclear moderate accumulation of Nrf2 is elevated in order to activate the CNC-bZIP factor and confer its potent resistance against acute drug toxicity [19]. Despite cytoprotective function of Nrf2 supported by graded expression of Keap1, a marked decrease of its levels to less than 50% results in increased mortality in a study of 2-year-old mice [20]. Overall, these facts demonstrate that the benefits of modest cytoprotective activation of Nrf2 against acute toxicity are hormetic, but its constitutive activation beyond a certain threshold is rather disadvantageous to long-term survival. The notion is supported by permanently hyperactive Nrf2 allowing for its transfiguration to act as an unrecognized mediator of oncogenesis and promote cancer cell survival in tumorigenesis [21, 22]. Hence, such distinct dual roles of Nrf2 (to become a good friend or dangerous foe) in tumor prevention and progression have led us to take careful account of its opposing potentials to implicate in cancer treatment.

In this present study, we first report a discovery of a naturally-occurring mutant of Keap1 (designated Keap1^ΔC^), which is expressed primarily in human highly-metastatic hepatoma MHCC97H cells, albeit it was widely expressed at much lower levels in all other cell lines examined. Keap1^ΔC^ is identified to arise from translation of an alternatively mRNA-spliced variant lacking the fourth and fifth exons of wild-type *Keap1* (but with no genomic deletion mutants). The resultant lack leads to a deletion of most of the Keap1 C-terminal domains required for binding Nrf2, to yield a dominant-negative mutant Keap1^ΔC^, acting as a competitor against inhibition of Nrf2-target genes by Keap1.

## RESULTS AND DISCUSSION

### Discovery of KEAP1^ΔC^ as an alternatively-spliced variant

The full-length protein of Keap1 was first identified to consist of 624 amino acids (aa) [11]. To date, there are at least 10 alternative splicing transcripts arising from the single *Keap1* gene (that contains 6 exons and 5 introns located in the chromosome 19p13.2) in GenBanks by NCBI (<u>Gene ID:9817</u>) and Ensembl (<u>ENSG00000079999</u>). Nevertheless, none of its protein isoforms have been reported so far. In addition, it is important to note that amino acid mutations of Keap1 were found to highly express in the human lung, breast and other somatic cancers [23–25]. These mutants were identified to exert significant disruptive effects on its dimerization *per se* and interaction with Nrf2, leading to the deteriorative activation of the CNC-bZIP factor.

Here, to confirm whether there exists a similar potential relationship between Keap1 and Nrf2, as accompanied by their mutants in hepatocarcinogenesis, we determined endogenous expression at their mRNA levels by real-time qPCR analysis of five hepatocellular carcinoma cell lines, followed by sequencing of their cDNAs. As shown in Figure 1A no changes in mRNA expression of Nrf2 were observed in these cancer cell lines, also with not any mutants examined (data not shown). By contrast, a considerable lower mRNA expression level of Keap1 was undetectable in HL-7702, a non-cancerous hepatocyte cell line, but not in other hepatocellular carcinoma cells (Fig. 1B). Interestingly, only the highly-metastatic hepatoma MHCC97H cells gave rise to double cDNA bands of Keap1, at much lower levels than those obtained from all other hepatocellular carcinoma cell lines, which just gave a single band of its full-length. Subsequent sequencing of the cDNA products from MHCC97H cells revealed that overlapped double peaks emerged from the 1326th nucleotide of the Keap1-coding region (Fig. 1C).

**Fig. 1.**
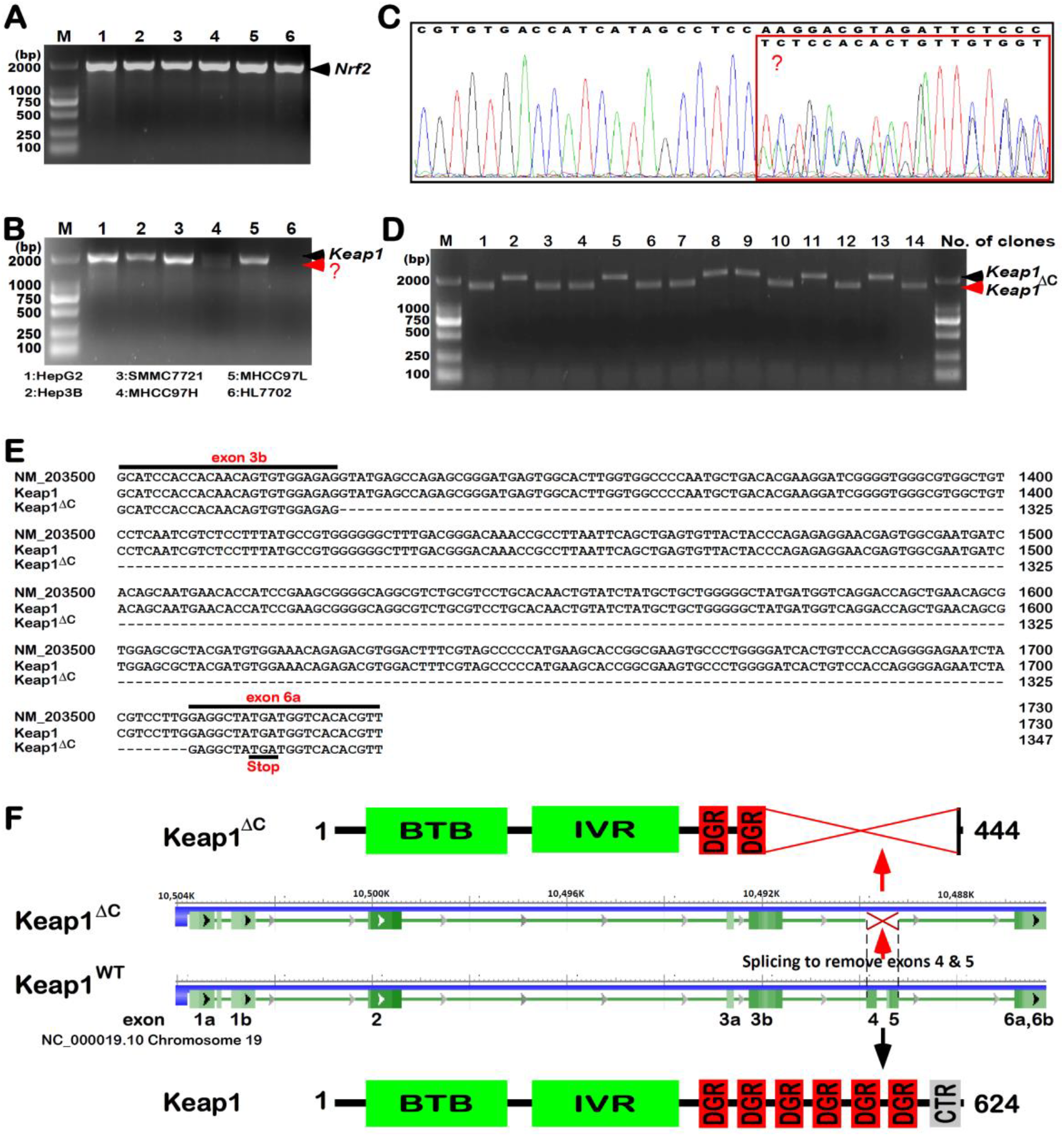
A novel discovery of the *Keap1*^*ΔC*^ mutant. Similar but different PCR products of *Nrf2* (A) and/or *Keap1* (B) were determined by 1% Agarose gel electrophoresis of samples from 6 cell lines as indicated on the bottom. (C) The products of *Keap1* PCR from MHCC97H cells were identified by sequencing and then analyzed by using the Chromas software. The overlapped peaks of the sequencing curves were placed in a red box. (D) Single colonies of *E. Coli* bringing a pcDNA3-based expression plasmid (in which the PCR products of *Keap1* from MHCC97H cells were inserted) were subjected to identification of inserted cDNA fragments by 1% Agarose gel electrophoresis. (E) A nucleotide sequence alignment of *Keap1* and *Keap1*^*ΔC*^ was listed, using the DNAMAN software, of which the GenBack Accession No. NM_203500 sequence served as a standard. (F) Shows schematic of distinct lengths of *Keap1* and *Keap1*^*ΔC*^ mRNAs, as well as structural domains of their proteins.

To clarify the overlapped double peaks of the above-described Keap1 cDNA sequences, they were subcloned into the pcDNA3 expression vector, before 14 clones were randomly selected and identified by PCR with a pair of Keap1 primers. The resulting electrophoresis of PCR products showed two types of bands with distinct sizes, though both were migrated closely to the standard 2000-bp DNA marker, on 1.0% Agarose gels (Fig. 1D). These two bands of Keap1 cDNAs were also determined by further sequencing of all selected clones. The result demonstrates that the relatively larger bands represent the wild-type full-length cDNA sequence of Keap1 without any mutation, whilst the shorter bands are owing to a loss of the entire fourth and fifth exons when compared with the intact wild-type (Fig. 1E). Together with not any of the genomic deletion mutants of the *Keap1* gene locus in these cells examined, these findings have led us to postulate there exists a novel alternatively-spliced variant of Keap1 transcript in MHCC97H cells. Further bioinformatic analysis of the spliced variant revealed a loss of the C-terminal 180-aa residues of Keap1 (called Keap1^ΔC^, Fig. 1F). This results in a constructive deletion mutant of most of the DGR domain and adjacent CTR region of Keap1 essential for its interaction with Nrf2 [23, 26]. Lastly, the nucleotide sequence of the variant Keap1^ΔC^ along with its amino acid sequence had been submitted to the GenBank developed by NCBI, with the accession No. MG770405 was granted.

### Differential expression of Keap1^ΔC^ in distinct cell lines

To determine whether Keap1^ΔC^ is also expressed in other cell lines rather than MHCC97H cells, two distinct pairs of primers targeting specifically for this variant or wild-type *Keap1* (Fig. 2A) were, according to their sequence similarity and difference, designed with the same upstream primers coupled with distinct downstream primers retaining the identical tetranucleotide (5’-CCTC-3’) at their 3’-ends, for a precision measure of real-time qPCR. Subsequent results illustrated that different products for Keap1 and Keap1^ΔC^ exhibited the same compatible melt peaks (Fig. 2B *left*), as accompanied by almost overlapped curves of their similar exponential amplification reactions (*right panel*). To validate the real-time qPCR with distinct pairs of primers, each of expression constructs for Keap1, Keap1^ΔC^ or empty pcDNA3 vector was transfected into HepG2 cells and then subjected to quantitative analysis.

**Fig. 2.**
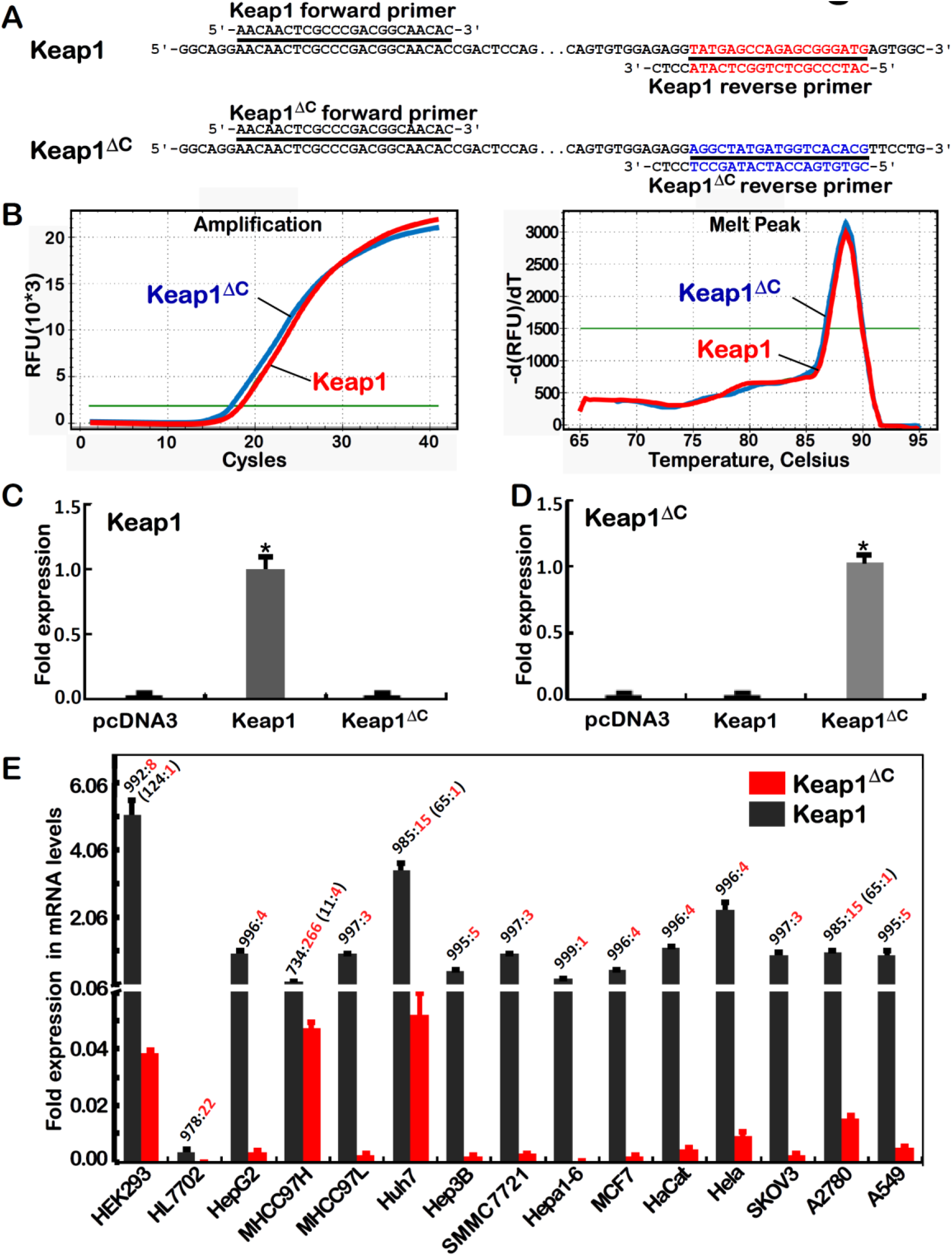
Distinctive expression of *Keap1*^*ΔC*^ in different cell lines. (A) Two pairs of primers were designed for specific binding *Keap1* and *Keap1*^*ΔC*^. their forward primers were identical, but their reverse primers had only 4 identical bases at the 3’ ends. (B) The *Keap1-* and *Keap1*^*ΔC*^-specific primers were identified by almost overlapped amplification curves (*left*) and melting curves (*right*) of real-time qPCR. (C,D) An expression construct for Keap1 and Keap1^ΔC^ was transfected into HepG2 cells. Then total RNAs were extracted and subjected to real-time qPCR to detect distinctive expression of *Keap1-* and *Keap1*^*ΔC*^-specific mRNAs (*p<0.01, n=3×3). (E) Different expression ratios of *Keap1* to *Keap1*^*ΔC*^ mRNA levels were determined by real-time qPCR analysis of 15 distinct cell lines as indicated.

As anticipated, the mRNA levels of ectopic *Keap1* and *Keap1*^*ΔC*^ were only detected by real-time qPCR with their respective specific primer pairs (Fig. 2, C & D), but not other products of their reciprocal chiasmata were measured. Further examinations revealed that endogenous *Keap1* and *Keap1*^*ΔC*^ mRNAs were widely but diversely expressed at varying extents in distinct types of 15 human somatic cancerous and non-cancerous cell lines (Fig. 2E), including 7 hepatoma-derived cell lines HepG2, MHCC97H, MHCC97L, Huh7, Hep3B, SMMC7721 and Hepa1-6, one lung cancer cell line A549, and other 4 female cancer cell lines MCF7 (from breast cancer), HeLa (from cervical carcinoma), SKOV3 and A2780 (the latter two from ovarian cancer), together with 3 non-cancerous cell lines of HL7702 (liver), Hacat (skin) and HEK293 (kidney). Overall, expression of *Keap1*^*ΔC*^ mRNA was at relatively higher levels determined primarily in MHCC97H, Huh7 and A2780 cells than other cell lines, but its *bona fide* abundances were also presented at considerable lower levels than the value of wild-type *Keap1* mRNA in all examined cell lines.

### An interference of Keap1^ΔC^ with Keap1 for its inhibition of Nrf2-target genes

To investigate whether inhibition of Nrf2 by Keap1 is interfered by distinct expression of Keap1^ΔC^ in four different cell lines, thus we examined basal protein levels of Keap1 and Nrf2, along with its downstream target genes *HO-1, GCLM* and *NQO1*. Western blotting showed a significant high abundance of Nrf2 only in MHCC97H cells, rather than other three cell lines determined (Fig. 3A1). This result is too coincidental to be just a coincidence, along with almost none of Keap1 detected in MHCC97H cells (Fig. 3A2), but as accompanied by relative higher levels of Keap1^ΔC^ (Fig. 2E). This finding indicates that the increase of Nrf2 should be attributable to remarked expression of Keap1^ΔC^. This notion is also supported by additional opposing fact that no increased, but considerable lower, levels of Nrf2 (Fig. 3A1) could result from almost absence of Keap1^ΔC^ in HL7702 cells having less expression of Keap1 (Figs. 3A2 and 2E). Conversely, significant higher abundances of Keap1 in MHCC97L and HepG2 cell lines (Fig. 3A2), along with very lower expression levels of Keap1^ΔC^ (Fig. 2E), led to marked decreases of Nrf2 to similar lower levels that measured from HL7702 cells (Fig. 3A1). Together, the non-cancerous HL7702 cells are allowed for maintaining proper constructive thresholds of Keap1 and Nrf2 to be expressed at controllable lower levels. In turn, it is inferable to exist a potential relevance of malignant hepatoma development to the constructive increases in basal Nrf2 or Keap1 proteins, which should also be affected by differential expression of Keap1^ΔC^. Further examinations of these cancerous and non-cancerous cells revealed a coupled positive and negative correlation of Nrf2 and Keap1, respectively, with basal protein expression levels of Nrf2-target genes HO-1 and GLCM, but not NQO1 (Fig. 3A3 to A5). Such an exception of NQO1 that was highly expressed in HL7702 and HepG2 cell lines (Fig. 3A5), though accompanied by lower levels of Nrf2, indicates that it may also be regulated by other CNC-bZIP family members (e.g. Nrf1) and other transcription factors (e.g. AP-1) [27, 28].

**Fig. 3.**
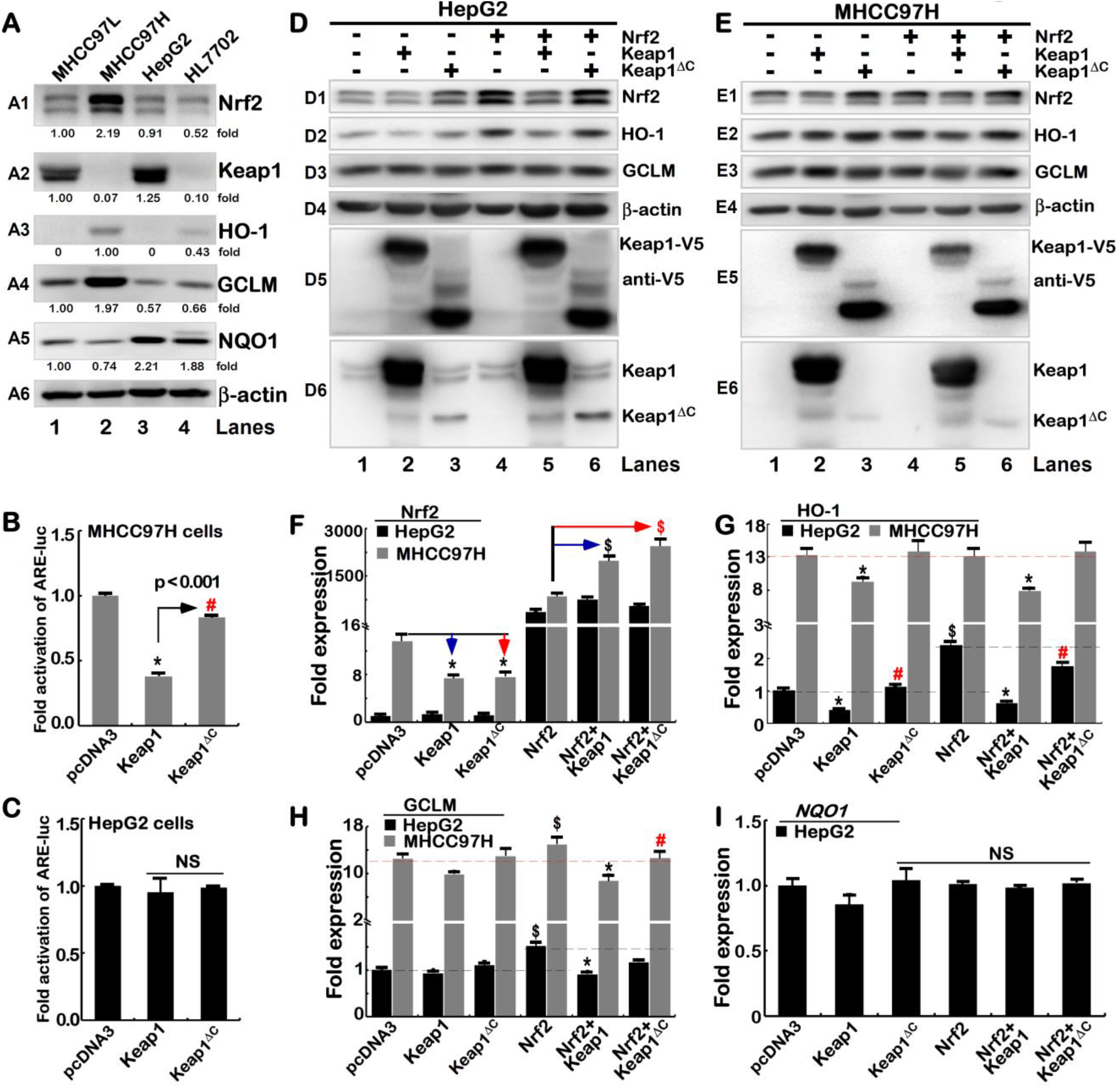
An effect of Keap1^ΔC^ is distinctive from Keap1 on Nrf2-target genes. (A) Different expression levels of endogenous proteins Keap1, Nrf2, HO-1, GCLM and NQO1 were determined by Western blotting of four distinct cell lines as indicated. β-actin served as a loading control. (B, C) An *ARE*-driven fluorescent reporter (pARE-Luc), an internal control pRL-TK together with an expression construct for Keap1 or Keap1^ΔC^, were co-transfected into MHCC97H (B) or HepG2 cells (C). After 24 h, the luciferase activity was assayed (*p<0.001, n=3×3; NS= not significant). Both HepG2 (D)and MHCC97H (E) cell lines were or were not co-transfected with expression plasmids for Keap1 or Keap1^ΔC^ alone or plus Nrf2. After 24 h the total lysates were subjected to Western blotting with distinct antibodies against Nrf2, HO-1, GCLM, β-actin, Keap1 and its V5 tag. (F to I) Similar transfected cells as described above were subjected to real-time qPCR analysis of *Nrf2, HO-1, GCLM* and *NQO1* at the mRNA expression levels. Significant increases (# or $p<0.01, n=3×3) and/or significant decreases(*p<0.01, n=3×3) were determined with error bars (S.E.).

To determine whether inhibition of Nrf2-mediated gene transcription by Keap1 is interfered by Keap1^ΔC^, both MHCC97H and HepG2 cell lines were co-transfected with a Keap1 or Keap1^ΔC^ expression construct together with an ARE-driven luciferase reporter. The results showed that over-expression of Keap1 caused a significant decrease in *ARE-Luc* reporter activity to ∼30% of control levels (Fig. 3B that were measured from transfection of MHCC97H cells with an empty pcDNA3 vector and the reporter). However, such a decreased transactivation activity of *ARE-Luc* reporter gene mediated by endogenous Nrf2 appeared to be, at least in part, disinhibited by endogenous Keap1^ΔC^ to a certain extent, albeit over-expression of Keap1^ΔC^ only resulted in slight repression of the reporter gene activity in the same MHCC97H cells (Fig. 3B). By contrast, Nrf2-transacting *ARE-Luc* reporter activity was almost unaffected by ectopic Keap1 or Keap1^ΔC^ in HepG2 cells (Fig. 3C). This is postulated to be owing to a considerable higher background of endogenous Keap1, so that Nrf2-targeted gene reporter may be not over-regulated by further over-expression of ectopic Keap1 or Keap1^ΔC^ in HepG2 cells.

To further determine the interfering role of Keap1^ΔC^ in the negative regulation of Nrf2 by Keap1, HepG2 and MHCC97H cell lines were transfected with a Keap1 or Keap1^ΔC^ expression construct alone or plus Nrf2 plasmids. As expected, Western blotting showed that endogenous protein levels of Nrf2 were obviously reduced (Fig. 3D1 & E1, *lanes 2 vs 1*) in both cell lines that had been allowed for further over-expression of ectopic Keap1 (Fig. 3D5, D6, E5 and E6). However, forced expression of ectopic Keap1^ΔC^ led to a significant augment in abundances of endogenous Nrf2 protein (*lanes 3 vs 1*). Further examinations also revealed that ectopic Keap1, but not Keap1^ΔC^, led to greater or less extents of distinct decreases in total levels of both ectopic and endogenous Nrf2 proteins (Fig. 3D1 and E1, *cf. lanes 5 & 6 with 4*). Moreover, variations in protein expression levels of different ARE-driven genes HO-1 (Fig. 3D2& E2), but not GCLM (Fig. 3D3 & E3), were correlated positively with changed abundances of Nrf2 *per se* in the presence or absence of ectopic Keap1 or Keap1^ΔC^, implying that HO-1 is an optimal marker of Nrf2-target genes.

Intriguingly, real-time qPCR showed that both endogenous and ectopic mRNA levels of *Nrf2* were almost not influenced by over-expression of either Keap1 or Keap1^ΔC^ in HepG2 cells, but the latter two regulators were required for striking suppression of endogenous *Nrf2*, but were also involved in significant promotion of its ectopic expression in MHCC97H (Fig. 3F), suggesting that *Nrf2* mRNA expression and/or its stabilization might be monitored by a proper status of Keap1 or Keap1^ΔC^. Importantly, further examinations revealed that both basal and Nrf2-mediated levels of *HO-1* mRNA were markedly repressed by ectopic Keap1, rather than Keap1^ΔC^, in HepG2 cells (Fig. 3G). By contrast, MHCC97H cells gave rise to an increased expression background of endogenous *HO-1* mRNA, but it was not further enhanced by Nrf2 over-expression, albeit its expression was obviously reduced by Keap1, but with almost none or less of inhibition by Keap1^ΔC^ (Fig. 3G). This implies that transcription of *HO-1* is regulated by Nrf2 in a steady-state system. In addition, basal *GCLM* and *NQO1* l mRNAs appeared to be unaffected by either Keap1 or Keap1^ΔC^ (Fig. 3H & I), but a marginal increase in Nrf2-induced *GCLM* rather than *NQO1* expression was diminished by Keap1. This was roughly not or less decreased by Keap1^ΔC^ in MHCC97H and HepG2 cells.

### Keap1^ΔC^ acts as a dominant-negative competitor against intact Keap1

To provide a better understanding of the axiomatic reason why Keap1^ΔC^ is enabled for its interference with Keap1, we conducted molecular modeling of two possible dimeric complexes of Keap1^ΔC^ (only retaining two Ketch motifs) with intact Keap1 (with all six Ketch motifs and adjacent C-terminal region, which are essential for direct association with Nrf2) (Fig. 4A). These models were based on a known crystal structure of Keap1 in complex with the N-terminal Neh2 region of Nrf2 (i.e. 3WN7 deposited in PDB)[15, 29, 30]. From formation of an invalid dimer of Keap1^ΔC^:Keap1 and Keap1^ΔC^:Keap1^ΔC^ (Fig. 4A *right panels*), it is therefore deduced that they do not only enable intact Keap1 to be simply consumed, but act as a potential dominant-negative competitor of Keap1 against its functional interaction with Nrf2.

**Fig. 4.**
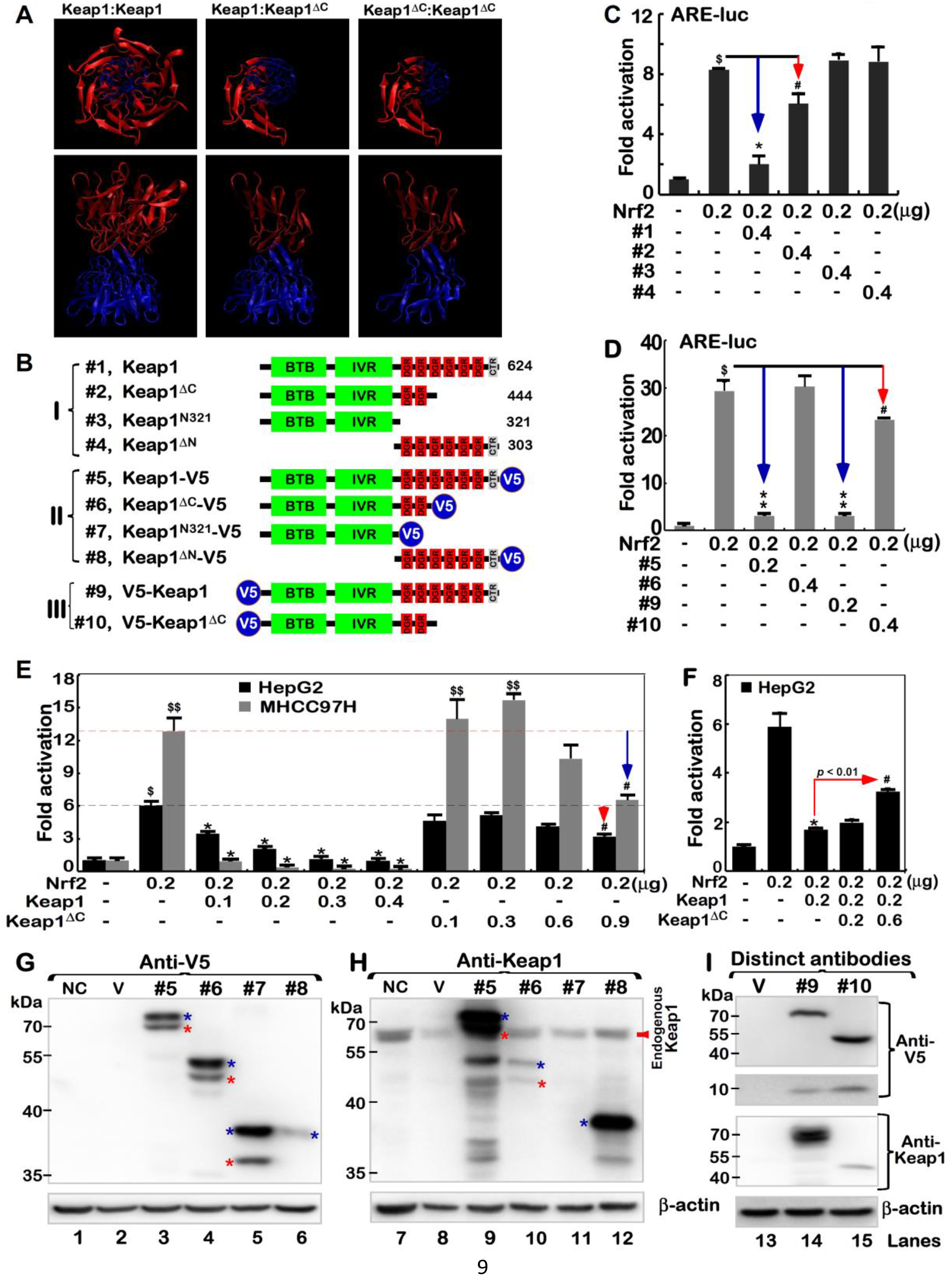
Keap1^ΔC^ competes against effects of Keap1 on Nrf2-mediated reporter gene. (A) Three distinct dimers of Keap1 and Keap1^ΔC^ with different views were simulated by the VMD1.9.3 software, based on a template of crystal structure of Keap1 (3WN7 deposited in PDB). (B) Shows schematic of the structural domains of Keap1, Keap1^ΔC^, and other mutants Keap1^N321^ and Keap1^ΔN^, some of which were tagged by the V5 ectope at the N-terminal or C-terminal ends. (C,D) HepG2 cells were co-transfected with an Nrf2 expression construct, and both reporters of *pARE-Luc* and *pRL-TK* (as an internal control), together with Keap1, Keap1^ΔC^ and other mutants as indicated. After 24 h, luciferase activity was measured with significant changes ($,p<0.001 compared to the normal control group; # p<0.01, * p<0.001 compared to the Nrf2-positive group; n=3×3). (E) HepG2 and MHCC97H cell lines that had been co-transfected with an Nrf2 expression construct and two reporters *pARE-Luc* and *pRL-TK*, in combination of distinct cDNA concentrations of Keap1 or Keap1^ΔC^, were subject to luciferase assays to determine ARE-driven reporter activity with significant increases ($, p<0.01; $$, p<0.001 compared to the normal control, n=3×3) or significant decreases (*, # p<0.01, compared to the positive Nrf2; n=3×3). (F) Besides an Nrf2 plasmid and two reporters *pARE-Luc* and *pRL-TK*, different concentrations of Keap1 and/or Keap1^ΔC^ alone or in combination were co-transfected into HepG2 cells. Then reporter gene activity was measured with significant changes (n=3×3; *p<0.001; #, p<0.01 compared to the Nrf2 positive group). (G-I) Expression constructs for Keap1-V5 (#5), Keap1^ΔC^-V5 (#6), Keap1^N321^-V5 (#7), Keap1^ΔN^-V5 (#8), V5-Keap1 (#9) or V5-Keap1^ΔC^ (#10), along with an empty pcDNA3 vector as the negative control (i.e. V), were transfected into HepG2 cells, respectively. The total lysates of either transfected or untransfected as a normal control (i.e. NC) cells were then analyzed by Western blotting with antibodies against Keap1 and its V5 tag.

To verify whether Keap1^ΔC^ has a marginal negativity to regulate Nrf2 through its residual DGR region, we created a series of expression constructs. Of note, the Keap1^N321^ mutant lacks the entire DGR domain, but retains its N-terminal 321-aa region, while Keap1^ΔN^ is an N-terminal deletion mutant, which comprises only 303 aa covering the entire DGR domain of Keap1 to its C-terminal end (Fig. 4B). Subsequently, ARE-driven luciferase reporter assays of HepG2 (Fig. 4C) and MHCC97H (Fig. 4D) cell lines demonstrated that Nrf2-mediated transactivation activity was significantly suppressed by Keap1, but was only marginally or not diminished by Keap1^ΔC^. The low-efficient effect of Keap1^ΔC^ appeared to be influenced by its C-terminal V5 tag (Fig.4D), albeit it is unknown of its functional folding. Further examinations revealed that Nrf2-target reporter activity was substantially repressed by over-expression of Keap1 in a dose-dependent manner (Fig. 4E). However, it is important to note that similar transactivation of this reporter gene mediated by Nrf2 was modestly induced by lower doses (cDNA) of Keap1^ΔC^ (without V5 tag), except diminishment by higher doses of Keap1^ΔC^ (Fig. 4E *right panel*). Particularly, inhibition of Nrf2 by Keap1 was blunted by Keap1^ΔC^, leading to marked de-repression of reporter gene activity (p< 0.01, Fig. 4F). The similar, but different, results were obtained from an additional aberrant-splicing mutant of Keap1 (with a loss of 458-624 aa) in the human prostate cancer DU-145 cells [31]. Collectively, these findings indicate a possibility that the Keap1^ΔC^ mutant may occupy competitively against intact Keap1 in the formation of an invalid dimer, as a consequence (Fig. 4A). Therefore, putative insufficient interaction of Keap1^ΔC^ with Nrf2 facilitates the CNC-bZIP factor to be released and translocated into the nucleus before transactivating target genes.

### Putative processing of Keap1 and Keap1^ΔC^ to yield distinct lengths of isoforms

The above-mentioned expression constructs (Fig. 4B) were also subject to further identification by Western blotting with distinct antibodies against the C-terminal residues 325-624 of Keap1 or the V5 ectope tagged to the N-terminal or C-terminal ends of this Kelch-like protein (Fig. 4 G and H). Curiously, the results revealed that Keap1 was likely subjected to the proteolytic processing of the protein within its N-terminal portein by a not-yet-identified protease, in order to yield two distinct lengths of isoforms between ∼76-kDa and 70-kDa (both C-terminally tagged by V5). Similar processing of the protein was also examined in additional two cases of Keap1^ΔC^ and Keap1^N321^ (Fig. 4G *lanes 3-5*). Besides the major ∼76-kDa full-length Keap1, its minor isoform of 70-kDa (which was migrated nearly to two close endogenous proteins, Fig. 4H) is inferable to be generated after possibly proteolytic removal of a small ∼8-kDa N-terminal polypeptide from intact Keap1. This is also further supported by additional observation that the small N-terminally V5-tagged polypeptide of ∼8-kDa was, by coincidence, yielded from the putative N-terminal proteolytic processing of Keap1 or Keap1^ΔC^ (Fig. 4I *lanes 14 & 15 in upper two panels*). Moreover, immunoblotting of Keap1^ΔN^ unraveled that additional post-synthetic processing of Keap1 might also occur within a region closer to its C-terminal end, such that the majority of the C-terminally-tagged V5 ectope of this mutant was truncated off the protein, albeit the remaining portion of Keap1^ΔN^ was still recognized by Keap1-specific antibody (Fig. 4G *lane 6 vs* Fig. 4H *lane 12*).

### An antagonist effect of Keap1^ΔC^ on Keap1-mediated turnover of Nrf2

To further explore the potential mechanism by which Keap1^ΔC^ de-represses the negative regulation of Nrf2 by Keap1, distinct settings of pulse-chase experiments were conducted in different cell lines. The COS-1 cells that had been transfected with an Nrf2-expression construct alone or in combination with an additional construct for Keap1 or Keap1^ΔC^, before being treated with cycloheximide (CHX, which inhibits biosynthesis of nascent proteins), were subjected to determination of whether Keap1^ΔC^ exerts an antagonist effect on Keap1-mediated turnover of Nrf2. As anticipated, over-expression of Keap1 caused a strikingly abrupt decrease in the abundance of Nrf2 to 16% of the basal expression levels in the non-Keap1-transfected cells (Fig. 5A). Conversely, a substantial increase to 1.67-fold amounts of Nrf2 resulted from co-expression of Keap1^ΔC^. Further examination revealed that over-expression of Keap1^ΔC^ rendered the half-life of Nrf2 turnover to be prolonged from 0.19 h (=11.4 min) to 0.28 h (=16.8 min) following treatment of cells with CHX (Fig. 5B). Even after 1-h treatment of cells with CHX, 20% of Nrf2 abundance was retained by Keap1^ΔC^, before being gradually decreased to its putative minimum of ∼7% by 2 h of the chemical treatment (Fig. 5A). However, just a 7% minimum of Nrf2 was also exhibited at time points of 0.31 h (=18.6 min) or 1.07 h (=64.2 min) respectively, following CHX treatment of cells that had been transfected with Keap1 or not (Fig. 5B).

**Fig. 5.**
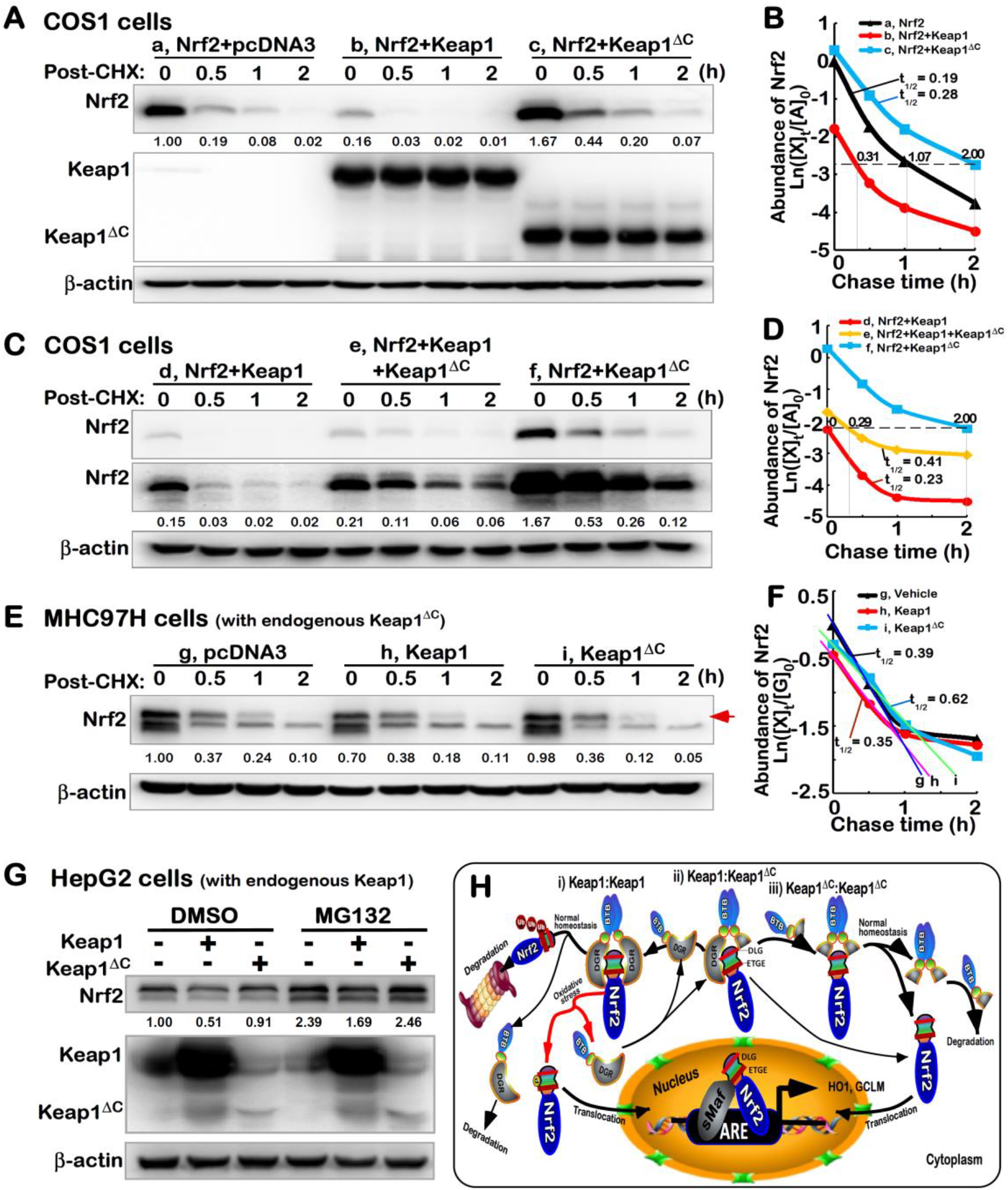
Keap1^ΔC^ has an antagonist effect on the Keap1-mediated turnover of Nrf2. (A to D) COS-1 cells were transfected with an Nrf2 expression plasmid alone or plus other plasmids V5-Keap1, V5-Keap1^ΔC^ or both, and then treated with 50μg/ml of CHX for distinct indicated lengths of time. The total lysates were determined by Western blotting with antibodies against Nrf2, Keap1, or its V5 tag, respectively. Subsequently, the intensity of blots was quantified and shown on *the bottom* (A & C). Of note, relative abundances of Nrf2 and its stability were determined with a changing half-life within distinct setting status, which were calculated and shown graphically (B & D). (E,F) MHCC97H and (G) HepG2 cells were transfected with an expression construct for V5-Keap1, V5-Keap1^ΔC^ or an empty plasmids, before being treated with 50μg/ml of CHX for indicated times or 10 μMol/L of MG132 for 4 h, prior to being harvested in a lysis buffer. Then, distinct abundances of endogenous Nrf2 proteins were examined by immunoblotting (E,G). The stability of endogenous Nrf2 protein was also determined with a varying half-life within distinct setting contexts, which was shown graphically (F). (H) A model is proposed to give a clear explanation of Keap1^ΔC^, acting as a dominant-negative competitor of Keap1. This is due to the fact that Keap1^ΔC^ can occupy the place in the formation of an invalid dimer with Keap1 or itself, no matter whether it only retains less or no ability to inhibit Nrf2. It is important to note that this mutant Keap1^ΔC^ has an antagonist effect on Keap1-mediated turnover of Nrf2 by proteasomal degradation pathway. In addition, upon dissociation of Nrf2 from Keap1, the CNC-bZIP factor will be allowed for spatio-temporal translocation into the nucleus before transactivating ARE-driven cytoprotective genes against oxidative stress or other biological stimuli.

The above-described data demonstrate that Keap1^ΔC^ is not a negative regulator of Nrf2, but conversely may act as an antagonist against endogenous Keap1, leading to disinhibition of keap1-mediated Nrf2 turnover. To address this hypothesis, COS-1 cells that had been co-transfected with expression constructs for Keap1, Keap1^ΔC^ or both together with Nrf2 plasmids were subjected to further pulse-chase experiments. As the time of CHX treatment was extended to 2 h, a fairly smooth tendency of decreases in the Nrf2 abundance from its maximum (1.67-fold) to its minimum (0.12-fold) was observed in cells co-expressing Keap1^ΔC^ and the CNC-bZIP protein (Figs. 5 C and D). By sharp contrast, the former minimum of Nrf2 was closer and even almost equal to additional starting level (0.15-fold) of this protein determined immediately before Keap1/Nrf2-co-expressing cells were treated with CHX (Fig. 5C *cf. the first lane with the last one*). However, the Keap1-led decrease of Nrf2 appeared to be partially mitigated, with a half-life slightly prolonged from 0.23 h to 0.41 h, by co-transfection of Keap1^ΔC^ (Figs. 5 C and D). This finding indicates that Keap1^ΔC^ exerts an antagonist effect on keap1 in monitoring the turnover of Nrf2 protein.

Further time-course analysis of CHX-treated MHCC97H cells revealed that endogenous Nrf2 protein levels were modestly reduced by over-expression of Keap1, rather than Keap1^ΔC^ (Fig. 5E), but its half-life was obviously extended by Keap1^ΔC^ from 0.39 h (=23.4 min) to 0.62 h (=37.2 min) following CHX treatment (Fig. 5F). Taken together with the data shown in Fig. 2E these suggest that endogenous Keap1^ΔC^ may compete Keap1, leading to de-repression of Keap1-mediated turnover of Nrf2 protein in MHCC97H cells. However, a marked increase in the endogenous CNC-bZIP protein was observed following treatment of HepG2 cells with proteasomal inhibitor MG132 (at 10 μmol/L) (Fig. 5G). Notably, over-expression of Keap1 rather than Keap1^ΔC^ caused significant decreases in baseline abundance and MG132-stimulated accumulation of endogenous Nrf2 protein in HepG2 cells. Taken together, these results demonstrate that Keap1, but not Keap1^ΔC^, is required for targeting Nrf2 to the proteasomal-mediated degradation pathway (Fig. 5H).

## CONCLUSION

As the most important inhibitory regulator of Nrf2, Keap1 plays a key role in monitoring the protein degradation of this CNC-bZIP factor and its subcellular locations, before regulating ARE-driven target genes [32, 33]. Here, we report a novel discovery of Keap1^ΔC^, as a naturally-occurring mutant of Keap1 with a constructive deletion of its C-terminal 180-aa residues essential for its directly binding to the Neh2 domain of Nrf2. The Keap1^ΔC^ mutant is generated from translation of the alternatively *Keap1* mRNA-spliced variant lacking both the fourth and fifth exons, but their coding sequences are retained in wild-type *Keap1* gene locus (i.e. with no genomic deletions). The variant Keap1^ΔC^ was determined primarily in the human highly-metastatic hepatoma MHCC97H cells, albeit is also widely expressed at very lower levels in all other cell lines examined. Further determination of Keap1^ΔC^ reveals that this mutant protein has a potent at forming an invalid dimer with Keap1 or itself, no matter whether it only retains less or no ability to inhibit Nrf2. As such being the case, the mutant Keap1^ΔC^ is also identified to act as a dominant-negative competitor of Keap1 against its negative regulation of Nrf2-target genes. This notion is also supported by additional fact that Keap1^ΔC^ exerts an antagonist effect on the Keap1-mediated turnover of Nrf2 protein. In addition, a line of new evidence has also been provided here, revealing that a putative N-terminal portion of Keap1 (and Keap1^ΔC^) can be also proteolytically truncated so as to yield its short isoforms, but this remains to be further determined.

## Materials and methods

### Chemicals, antibodies and other reagents

All chemicals were of the highest quality commercially available. The 26S proteasome inhibitor (MG132, bortezomib) were purchased from Sigma-Aldrich (St Louis, MO, USA). The primary antibodies such as V5 (Invitrogen, Shanghai), KEAP1 (D154142, Sangon Biotech), as well as Nrf2 (ab62352) and its target HO1 (ab52947), GCLM (ab126704) and NQO1 (ab80588) (all the latter four antibodies from Abcam, China) were herein employed. In addition to another first antibody against β-actin, various secondary antibodies were obtained from ZSGB-BIO (Beijing, China).

### Cell lines

All these cell lines HEK293, HL7702, HepG2, MHCC97H, MHCC97L, Huh7, Hep3B, SMMC7721, Hepa1-6, MCF7, Hacat, Hela, SKOV3, A2780 and A549 used in this study were purchased from the Cell Bank of Type Culture Collection of Chinese Academy of Sciences (Shanghai, China) and maintained according to the supplier’s instructions.

### Expression constructs and transfection

The human full-length Nrf2, Keap1 and Keap1^ΔC^ cDNA sequences were subcloned into a pcDNA3 vector, respectively. Deletion mutants of Keap1 were created by inserting appropriate PCR-amplified cDNA fragments into the above-described vector. Related primers were listed in Table 1. These mutants included Keap1^ΔN^ (with a deletion of the N-terminal 321-aa from Keap1), Keap1^N321^ (only retaining the N-terminal 321-aa region). In addition, they were N-terminally or C-terminally tagged by the V5 ectope to yield distinct strategic expression constructs. Following all these constructs were verified by DNA sequencing, they were transfected into experimental cells (2.5×10^5^), which had been allowed for growth to a confluence of 80% in 35-mm culture dishes. The transfection was carried out for 15-min in a mixture of indicated constructs (2 μg/ml) with Lipofectamine 3000 (Invitrogen, USA), and then allowed for a recovery from transfection before being experimented.

**Table 1.**
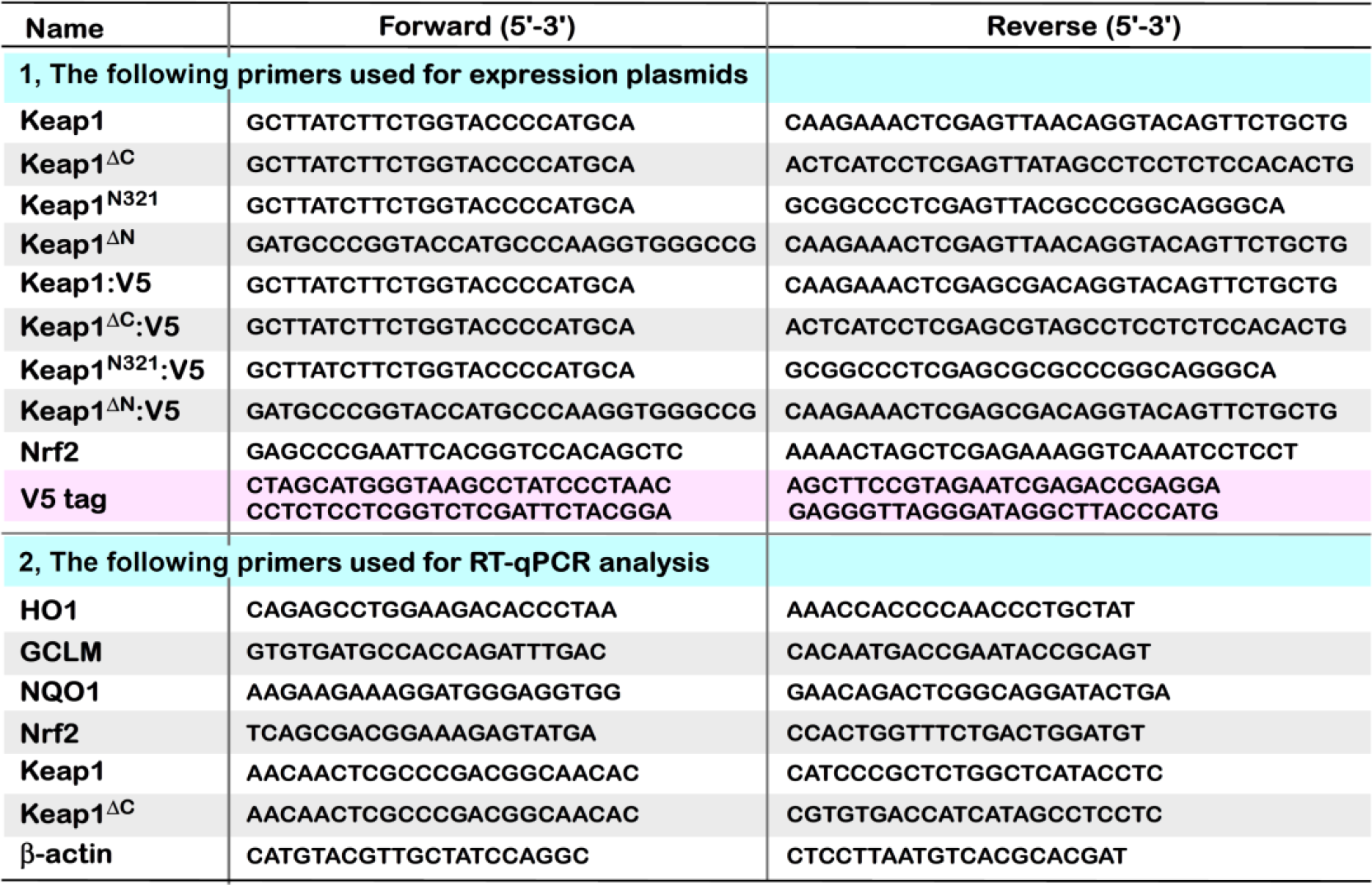
Distinct pairs of primers used for expression constructs and PT-qPCR analysis.

### Luciferase reporter assay

A *pARE-Luc* reporter plasmid [34], which was used for a measure of the transactivation activity of ARE-driven gene mediated by Nrf2, and pRL-TK, which is an internal control of Renilla luciferase reporter, together with indicated expression constructs were co-transfected into HepG2 or MHCC97H cells. At about 24 h after transfection, the luciferase reporter activity was measured by using dual luciferase reporter assay (Promega, USA).

### Real-time quantitative PCR

Experimental cells that had been transfected or not been transfected with the indicated plasmids were subjected to isolation of total RNAs by using the RNAsimple Kit (Tiangen Biotech CO., Beijing). Subsequently, 500 ng of total RNAs was added in a reverse-transcriptase reaction to generate the first strand of cDNA (with Revert Aid First Strand Synthesis Kit from Thermo). The synthesized cDNA was served as the template for qPCR, in the GoTaq® qPCR Master Mix (from Promega), before being deactivated at 95°C for 10 min, and amplified by 40 reaction cycles of 15 s at 95°C and 30 s at 60°C. The final melting curve was validated to examine the amplification quality, whereas expression of mRNA for β-actin was viewed as an internal standard control.

### Western blotting

Experimental cells were or were not transfected with indicated constructs for ∼8 h, and then allowed for a recovery culture for 24 h in a fresh DMEM containing 25 mmol/L high glucose. The cells were treated with indicated chemicals for 30 min to 4 h before being harvested in denatured lysis buffer (0.5% SDS, 0.04 mol/L DTT, pH 7.5) that contained the protease inhibitor EASYpacks (1 tablet/10 mL, Roche, Germany). The lysates were denatured immediately at 100°C for 10 min, sonicated sufficiently, and diluted in 3× loading buffer (187.5 mmol/L Tris-HCl, pH 6.8, 6% SDS, 30% Glycerol, 150 mmol/L DTT, 0.3% Bromphenol Blue) at 100°C for 5 min. Subsequently, equal amounts of protein extracts were subjected to separation by SDS-PAGE containing 4-15% polyacrylamide, and visualization by Western blotting with distinct antibodies. On some occasions, the blotted membranes were stripped for 30 min and then re-probed with additional primary antibodies. β-actin served as an internal control to verify equal loading of proteins in each of electrophoretic wells.

### Statistical analysis

Significant differences in the ARE-driven reporter activity mediated by Nrf2 and its target gene expression levels were **s**tatistically determined using either the Student’s t-test or Multiple Analysis of Variations (MANOVA). The data are shown as a fold change (mean ± S.D.), each of which represents at least 3 independent experiments that were each performed triplicate.

## Author contributions

L.Q. designed and performed most experiments, and prepared drafts of the paper with figures. Y.X. had done part of the work as shown in Figure 5. M.W. helped with data statistics and calculations. Y.P.Z. helped L.Q. to do with reporter assays. Y.Z. designed this study, supervised all the experiments, analyzed all the data, helped to prepare all figures, wrote and revised the paper.

## Acknowledgments

The study was supported by the National Natural Science Foundation of China (key programs 91129703, 91429305 and project 31270879) awarded to Prof. Yiguo Zhang (University of Chongqing, China), and also in part funded by Chongqing University postgraduates’ innovation project (No. CYB15024) funded to Mr. Qiu Lu. We are greatly thankful to Zhengwen Zhang (working at the Institute of Neuroscience and Psychology, School of Life Sciences, University of Glasgow, 42 Western Common Road, G22 5PQ, Glasgow, Scotland, United Kingdom) for his assistance to help the English correction of the paper.

